# Investigating the role of insulin in increased adiposity: Bi-directional Mendelian randomization study

**DOI:** 10.1101/155739

**Authors:** RC Richmond, KH Wade, L Corbin, J Bowden, G Hemani, NJ Timpson, G Davey Smith

## Abstract

Insulin may serve as a key causal agent which regulates fat accumulation in the body. Here we assessed the causal relationship between fasting insulin and adiposity using publicly-available results from two large-scale genome-wide association studies for body mass index and fasting insulin levels in a two-sample, bidirectional Mendelian Randomized approach. This approach is only valid on the condition that the two instruments are independent of one another. In analysis excluding overlapping loci, there was an increase of 0.20 (0.17, 0.23) log pmol/L fasting insulin per SD increase in BMI (P= 2.80 x 10^−36^), while there was a null effect of fasting insulin on BMI, with a 0.01 (−0.39, 0.38) SD decrease in BMI per log pmol/L increase in fasting insulin (P= 0.98). Furthermore, a high degree of heterogeneity in the causal estimates was obtained from the insulin-related variants, which may be attributed to varying mechanisms of action of the insulin-associated variants. Results were largely consistent when an Egger regression technique and weighted median and mode estimators were applied. Findings suggest that the positive correlation between adiposity and fasting insulin levels are at least in part explained by the causal effect of adiposity on increasing insulin, rather than vice versa.

## Introduction

The energy balance model of obesity based on the First Law of Thermodynamics states that energy intake minus energy expenditure is equal to change in stored energy (i.e., weight gain or weight loss)^1^. Opponents of the energy balance hypothesis criticize its circular reasoning which, they state, does not separate out cause and consequence of overeating and weight gain and therefore does not provide a primary driver for the obesity epidemic.^2,3^ Recently there has been a revival of a theory about the obesity epidemic which provides an alternative mode of thinking to the prevailing hypothesis. This alternative hypothesis highlights the role of carbohydrate (and in particular, refined sugars such as sucrose and fructose) as being responsible for increasing rates of obesity and identifies insulin as the key causal agent that is driven by the carbohydrate content of diet and which regulates fat accumulation in the body.^2^

The “insulin-carbohydrate” hypothesis views obesity as a hormonal, regulatory disorder and, in doing so, identifies insulin as a causal agent which may be directly targeted to reduce adiposity. Thus, this hypothesis provides impetus for the development of diet regimens that aim to lower intake of carbohydrate and glycaemic load.^3^ However, although observationally there is a strong association between levels of insulin and measures of adiposity, causal mechanisms (and indeed the direction of causation) still remain uncertain^4^. As a result, critics argue that this alternative hypothesis provides unwarranted support for fad diets proposed to lower levels of obesity^5^·^6^.

Randomised controlled trials (RCTs) examining the effectiveness of low sugar diets have generally been limited by short trial duration, a lack of compliance and little differentiation between treatment groups. Where an effect of increased sugar intake on weight change has been observed, data suggest that this is mediated through changes in total energy intake.^7^ Meanwhile, other trials suggest that insulin therapy may have a direct effect on weight gain and proposed mechanisms include the anabolic effects of high-dose insulin and appetite increases.^8^ However, such trials have generally been performed in groups of diabetic patients and so findings may not be generalisable to the general population, where research into the role of hyperinsulinemia on obesity in the absence of insulin resistance and glucose intolerance is required.

It may be possible to investigate causality in the association between insulin and adiposity using alternative methods of causal inference. Mendelian randomization (MR) is an approach that uses genetic variants robustly associated with modifiable exposures to infer causality.^9,10^ Such variants are not theoretically or empirically related to potential confounding factors or influenced by the development of the outcome. Therefore, they are not subject to confounding or reverse causation, allowing for the unbiased estimation of causal effects. The MR design is analogous to an RCT, where study participants are randomly allocated to one or another treatment, avoiding potential confounding between treatment and outcome.^11^

Statistical methodology for MR has been extensively developed since it was initially proposed^10^ and developments include: the use of multiple genetic variants to improve the precision with which causal effects may be estimated^12,13^; the application of two-sample MR based on summary statistics from two independent studies for genotype-exposure and genotype-outcome assessment^13–15^; bi-directional MR, which may be used in situations where the direction of causality in an observed association is uncertain^16^; and the application of various sensitivity analyses to test the assumptions of the MR approach, particularly pertaining to an appraisal of invalid instruments, which may bias causal estimates^15,17,18^.

We aimed to investigate the causal relationship between fasting insulin and adiposity using publicly available results from two large-scale genome-wide association studies (GWAS) for body mass index (BMI)^19^ and fasting insulin levels^20^ in a two-sample, bidirectional MR approach.

## Methods

### Defining genetic instruments

For BMI, a genetic instrument was constructed using 97 BMI-associated SNPs and their effect sizes (and associated standard errors) as estimated in a large-scale multi-ethnic meta-analysis of Genome wide Association Studies conducted by the Genetic Investigation of Anthropometric Traits (GIANT) consortium (n = 339,224)^19^. To generate the corresponding SNP-outcome (i.e., insulin) association, we took effect estimates and standard errors from publicly available results of a GWAS meta-analysis conducted by the Meta-Analysis of Glucose and Insulin-related traits Consortium (MAGIC) (n = 133,010)^20^.

For fasting insulin, a genetic instrument was constructed using 14 fasting insulin-associated SNPs and their effect sizes (and associated standard errors) as estimated in a large-scale meta-analysis of GWAS in the MAGIC consortium^20^. Although 19 SNPs were identified as being robustly associated with insulin in this meta-analysis, an additional 5 SNPs were identified in a GWAS of insulin only when adjusted for BMI; therefore, these SNPs were excluded due to potential bias induced in the genetic associations in this context^21^. To generate the corresponding SNP-outcome association (i.e., BMI), we took effect estimates and standard errors from a GWAS meta-analysis conducted by the GIANT consortium^19^.

Where the exposure SNPs were not available in the outcome data, we identified proxy SNPs in linkage disequilibrium (*r*^2^ > 0.8 and within 250kb of the target SNP) to use in analyses. Given an absence of proxy SNPs for 7 of the BMI SNPs in the fasting insulin GWAS, we repeated analyses using summary data from a previous GWAS effort by MAGIC^22^, consisting of data obtained in up to 46,186 non-diabetic participants. GWAS summary results available for this study consisted of 1,860,357 SNPs directly genotyped or imputed from HapMap CEU sample data, compared with 64,436 SNPs directly genotyped on the MetaboChip array, which were available in the larger study used in our main analysis. All 97 BMI SNPs were present in this GWAS summary dataset and therefore there was no need to identify proxy SNPs.

### Statistical analyses

All analyses were performed using the MR-Base “TwoSampleMR” package^23^ in R version 3.2.2. We harmonized the SNP-exposure and SNP-outcome associations using the MR-Base “harmonise_data” function^23^ to ensure that the associations obtained from the exposure and outcome GWAS summary-level data were coded relative to the same effect allele of each SNP. All harmonized SNP-exposure and SNP-outcome associations were combined using the inverse-variance weighted (IVW) method^17^. This produces a causal estimate of the exposure-outcome association, which is equal to the coefficient from a weighted regression of the SNP-outcome on the SNP-exposure association estimates, where weights are the inverse of the precision of the SNP-outcome coefficients and the intercept is constrained to zero.

## Sensitivity analyses

### Investigating potential pleiotropy and prevailing direction of effect

The existence of directional pleiotropy, where a genetic instrument has an effect on an outcome independent of its effect on the exposure, is a particular problem for MR, especially when using multiple genetic variants where the function of variants identified in GAWS is unknown.

One indication of pleiotropy is a high degree of heterogeneity between causal estimates of the individual SNPs combined in the IVW method. In the meta-analysis of the individual SNP estimates, we therefore calculated Cochran’s Q-statistic to estimate the degree of heterogeneity (p[het]) and Cook’s distance^24^ as a measure of the aggregate impact of each SNP in the model. We also applied the MR-Egger regression technique^17^, which was used to test overall directional pleiotropy and provide a valid causal estimate, taking into account the presence of pleiotropy. Violation of the no measurement error (NOME) assumption of the SNP-exposure estimates was addressed using simulation extrapolation (SIMEX) to adjust the MR-Egger estimate for regression dilution ^25^. Further to this, we applied the weighted median method, which provides a valid causal estimate under the assumption that up to 50% of the weight in the analysis stems from instrumental variables are invalid^18^, and the weighted mode method, which provides valid estimates when the largest number of similar causal effect estimates comes from valid instruments, even if the majority of instruments are invalid^26^.

An important caveat relevant to bidirectional MR, is that these methods are only valid on the condition that the two instruments are independent of one another (i.e., that there is no overlap or linkage disequilibrium between genetic variants included in the construction of the instruments used for the two traits being investigated)^10^. For example, it is possible that an instrument for insulin may be contaminated by genetic variants directly associated with BMI if there is a causal effect of BMI on insulin. This leads to a violation of the InSIDE (Instrument Strength Independent of Direct Effect) assumption of MR-Egger, which states that the direct pleiotropic effect of the genetic variants on the outcome must be independent of the instrument strength (i.e., the association between the instrument and exposure)^17^. Therefore, we investigated whether any of the 97 BMI-associated variants and the 14 insulin-associated variants were identical or strongly correlated and repeated the analyses following the exclusion of such SNPs.

As a final sensitivity analysis, we used the Steiger test to provide evidence for the prevailing causal direction, based on the estimated variance explained by the SNPs in the exposure and the outcome^27,28^.

### Investigating insulin secretion vs insulin resistance

Previous studies indicate that genetic variants associated with fasting insulin are involved in different pathways of insulin action, specifically insulin resistance and secretion^20,29^. Of the 14 SNPs associated with fasting insulin that were used in our main analysis, 6 were previously identified as being directly involved in insulin resistance as they showed associations with low HDL and higher triglycerides^20^, hallmarks of common insulin resistance **(Supplementary Table 1).^29^** To investigate the relevance of these pathways within the context of BMI, we stratified our SNP set into those deemed to be “insulin resistance” SNPs (N=6) and those that were not (N=8) and compared causal estimates with BMI obtained between these strata.

In an extension of this, we used a further list of SNPs^29^, which included an additional 4 variants associated with insulin resistance (totaling 10 SNPs) and 21 variants associated more specifically with insulin secretion **(Supplementary Table 2),** and then compared the causal effect of fasting insulin on BMI between these strata.

## Results

Using the available 90/97 SNPs robustly associated with BMI that were presentin the MAGIC GW AS summary results (total r^2^ = 2.5%; F-statistic = 95.6), there was a 0.21 (95% CI 0.17, 0.24) log pmol/L increase in fasting insulin per SD (4.5kg/m^2^) increase in BMI (P=1.57×l0'^32^), although there was some evidence of heterogeneity for the individual causal estimates obtained for each variant (P(het) = 7.07xl0^13^) **(Table 1, Figure 2),**

**Table 1.** Results of Mendelian randomization analysis with body mass index as the exposure and insulin as the outcome

| N SNPs | Method | Coefficient | Estimate<sup>±</sup> | 95% CI | p-value |
| --- | --- | --- | --- | --- | --- |
| 90 | IVW | Slope ($\beta$) | 0.21 | 0.17, 0.24 | 1.57x10<sup>-32</sup> |
| 90 | MR-Egger | Slope ($\beta$) | 0.19 | 0.10, 0.27 | 3.40x10<sup>-5</sup> |
| 90 | MR-Egger | Intercept ($\alpha$) | 0.001 | -0.002, 0.003 | 0.62 |
| 90 | SIMEX* | Slope ($\beta$) | 0.21 | 0.11, 0.30 | 3.47x10<sup>-5</sup> |
| 90 | Weighted Median | Slope ($\beta$) | 0.23 | 0.19, 0.26 | 1.57x10<sup>-35</sup> |
| 90 | Weighted Mode | Slope ($\beta$) | 0.24 | 0.18, 0.29 | 5.31x10<sup>-14</sup> |

\*Weighted I-squared=0.88
<sup>±</sup> log pmol/L increase in fasting insulin per SD (4.5kg/m<sup>2</sup>) increase in BMI

**Figure 1.**
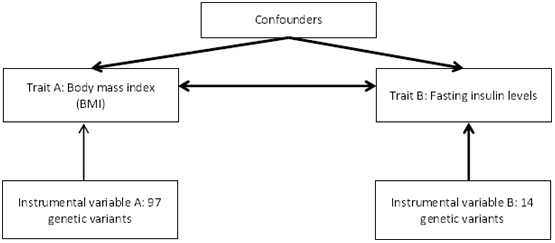
Schematic representation of bidirectional Mendelian randomization of BMI and insulin

**Figure 2.**
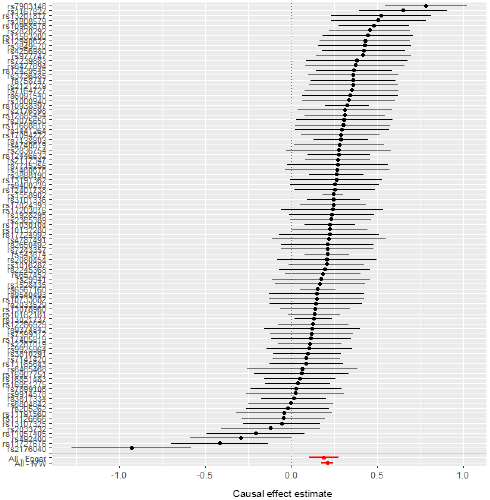
Mendelian randomization analysis estímate of the association of genetically-elevated body mass index on fasting insulin levels using 90 SNPs robustly associated with BMI as instrumental variables

While the results of the MR analysis provided suggestive evidence for a causal effect of adiposity on increasing insulin levels using data from the most recent GIANT^19^ and MAGIC^20^ GWAS efforts for BMI and fasting insulin, respectively, we repeated analyses using summary data on all 97 SNPs robustly associated with BMI, which were present in the previous MAGIC GWAS effort^22^, consisting of data obtained in up to 46,186 nondiabetic participants. The IVW method gave a causal estimate similar to that previously obtained in the main analyses (0.18 (95% CI 0.14, 0.21) log pmol/L increase in fasting insulin per SD increase in BMI (P=2.02×l0^−20^)). There was less evidence of heterogeneity for the individual causal estimates obtained for each variant (P(het) = 0.05).

In the reverse direction, using 14 SNPs robustly associated with fasting insulin that were present in the GIANT GWAS summary results (r^2^ = 0.4%; F-statistic = 35.0), there was a 0.58 (95% CI −0.20,1.36) SD increase in BMI per log pmol/L increase in fasting insulin (P=0.14) **(Table 2, Figure 3).** However, there was evidence of heterogeneity for the individual causal estimates obtained for each fasting insulin-associated variant (P(het) =1.83 × 10^−146^).

**Table 2 -.** Results of Mendelian randomization analysis with insulin as the exposure and body mass index as the outcome

| N SNPs | Method | Coefficient | Estimate<sup>±</sup> | 95% CI | p-value |
| --- | --- | --- | --- | --- | --- |
| 14 | IVW | Slope ($\beta$) | 0.58 | -0.20, 1.36 | 0.14 |
| 14 | MR-Egger | Slope ($\beta$) | 3.76 | -0.48, 8.00 | 0.11 |
| 14 | MR-Egger | Intercept ($\alpha$) | -0.05 | -0.13, 0.02 | 0.16 |
| 14 | SIMEX* | Slope ($\beta$) | 5.93 | -0.71, 12.57 | 0.08 |
| 14 | Weighted Median | Slope ($\beta$) | 0.16 | -0.06, 0.38 | 0.14 |
| 14 | Weighted Mode | Slope ($\beta$) | 0.13 | -0.13, 0.39 | 0.36 |

\*Weighted I-squared=0.25
<sup>±</sup>SD increase in BMI per log pmol/L increase in fasting insulin

**Figure 3.**
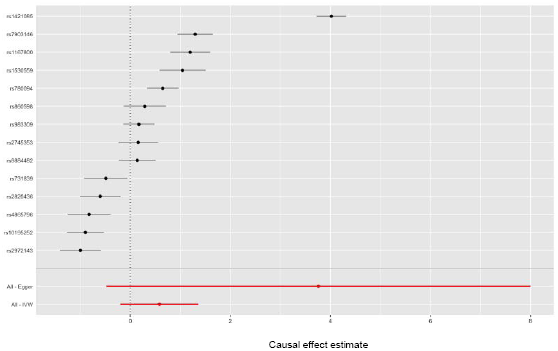
Mendelian randomization analysis estimate of the association of genetically-elevated fasting insulin levels on body mass index using 14 SNPs robustly associated with BMI as instrumental variables

## Sensitivity analysis

### Investigating potential pleiotropy and prevailing direction of effect

All approaches provided evidence of a positive causal effect of BMI on insulin, with the causal estimate from both the MR-Egger, weighted median and weighted mode being very similar to that obtained by the IVW method (**Table 1, Supplementary Figure 1a**). When plotting the instrument strength (i.e., the genetic associations with BMI) against the individual IV estimates in a funnel plot, there was a degree of asymmetry (**Supplementary Figure 1b**), suggesting some level of directional pleiotropy. However, this was largely driven by one variant in the *HHIP* gene region and the MR-Egger intercept estimate of 0.001 (95% CI −0.002, 0.003; P = 0.62) was consistent with the null. Calculation of Cook’s distance showed four variants (including the *HHIP* variant) to have a disproportionate level of influence on the model compared to the other variants in the set (**Figure 4a**). Three out of four of these variants were found to be identical or in strong LD with genetic instruments for fasting insulin (**Table 3**).

**Figure 4 -.**
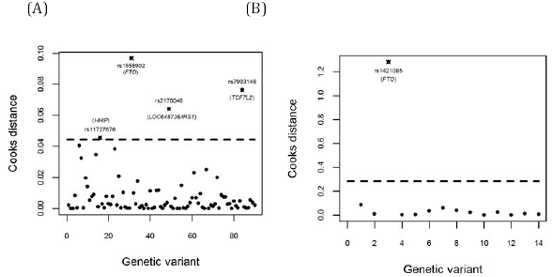
Cook’s distance applied to the IVW analysis to assess outliers (A) for effect of BMI on fasting insulin (B) for effect of fasting insulin on BMI Figure 5 – Genetic associations between BMI and fasting insulin stratified by relationship with insulin resistance

**Table 3 -.** Overlapping loci between BMI and insulin genetic instrument lists

| Locus | BMI instrument SNP | Insulin instrument SNP | Linkage disequilibrium (r<sup>2</sup>) |
| --- | --- | --- | --- |
| FTO | rs1558902 | rs1421085 | 1.000 |
| TCF7L2 | rs7903146 | rs7903146 | - |
| HIP1 | rs1167827 | rs1167800 | 0.680 |
| LOC646736/IRS1 | rs2176040 | rs2972143 | 0.963 |

All approaches provided suggestive evidence of a positive causal effect of insulin on BMI, with the causal estimate from MR-Egger being larger than the weighted median, weighted mode and IVW estimates (3.76 SD (95% CI −0.48, 8.00; P = 0.11), although with wide confidence intervals (**Table 2, Supplementary Figure 2a**). The asymmetry shown in the funnel plot of the instrument strength (i.e., the genetic associations with insulin) and the IV estimates (**Supplementary Figure 2b**) provided some evidence of negative directional pleiotropy, although the MR Egger regression was consistent with the null (intercept = −0.05 (95% CI −0.14, 0.02); P = 0.16). Calculation of Cook’s distance showed one variant, in the *FTO* gene region, to have a disproportionate level of influence on the model compared to the other variants in the set (**Figure 4b**). This variant is in strong LD with another SNP used as a genetic instrument for body mass index (**Table 3**).

Of the insulin-associated SNPs used in this analysis, the *FTO* variant was most likely to have an effect on fasting insulin entirely mediated through BMI^20^. In line with this and as anticipated, there was evidence for a strong positive pleiotropic effect of the *FTO* variant on BMI (**Supplementary Figure 2a**). However, as the MR-Egger approach assumes that causal estimates for the stronger genetic variants should be closer to the true causal effect, this approach treats the effect estimates obtained from variants such as *FTO* as the least invalid^17^. As such, the MR-Egger estimate indicates directional pleiotropy in the negative direction when, in actual fact, the pleiotropic effect of *FTO* may induce a spurious positive effect of insulin on BMI in both the IVW and MR-Egger analyses. We therefore repeated the analyses, excluding *FTO* along with any other variants that were found to overlap the list of genetic variants for both BMI and insulin.

In analysis excluding overlapping loci, similar estimates for the causal effect of BMI on insulin to that previously identified were obtained for all three approaches, with a 0.20 (0.17, 0.23) log pmol/L increase in fasting insulin per SD increase in BMI (P= 2.80 × 10^−36^) in the IVW analysis and a 0.12 (0.03, 0.21) log pmol/1 increase in fasting insulin (P=0.01) in the MR-Egger analysis (**Supplementary Table 3, Supplementary Figure 3**). The intercept of the MR-Egger regression was consistent with the null (intercept = 0.002 (−0.0001, 0.004), P = 0.06), suggesting no strong directional pleiotropic effect However, evidence for heterogeneity of the individual causal estimates obtained for each BMI-associated variant on fasting insulin remained (P(het) = 8.70 × 10^−5^).

Using summary data on the full 97 SNPs from the previous MAGIC GW AS effort^22^, effect estimates from MR-Egger, weighted median and weighted mode approaches were also consistent with this analysis, and the MR-Egger intercept (0.0003 (95% CI −0.003, 0.002)) indicated no directional pleiotropy (**Supplementary Table 4**). Furthermore, results were largely unchanged after removing overlapping SNPs associated with fasting insulin (IVW estimate = 0.17 (95% CI 0.13, 0.21; P=4.47×l0^−19^, P(het)=0.24) (**Supplementary Table 5**).

With the exception of the weighted median and weighted mode approaches, estimates for the causal effect of insulin on BMI attenuated with the exclusion of these overlapping variants, with a 0.01 (−0.39, 0.38) SD decrease in BMI per log pmol/L increase in fasting insulin (P= 0.98) in the IVW analysis (**Supplementary Table 6, Supplementary Figure 4**), The intercept of the MR-Egger regression was negative, although with a smaller magnitude and wide confidence intervals (intercept = −0.02 (95% CI −0.06, 0.02), P = 0.39), providing no strong evidence of directional pleiotropy. Furthermore, calculation of Cook’s distance showed no outliers (**Supplementary Figure 5a**). However, there was evidence for heterogeneity of the individual causal estimates obtained for each fasting insulin-associated variant on BMI (P(het) = 1.58 x 10^−13^)

There was evidence for heterogeneity in the individual SNP estimates in all of the stratified analyses; therefore, it remains possible that there is violation of the InSIDE assumption in these analyses (for example if some of the SNPs being used as instruments for insulin resistance or insulin secretion have a direct effect on BMI), However, the Steiger test provided evidence against this in most analyses, where the tested direction of effect was found to be the prevailing causal direction (**Supplementary Table 7**),

### Investigating insulin secretion vs insulin resistance

One explanation for the level of heterogeneity observed in the individual causal estimates for each fasting insulin-associated variant on BMI is the action of the genetic variants in either insulin secretion or insulin resistance pathways.^20^ We therefore performed stratified analysis to establish causal effects for 6 “insulin resistance” SNPs (**Supplementary Table 8**) compared with the remaining 8 variants, based on findings from a previous study^20^.

The causal effect of fasting insulin on BMI was inverse using the SNPs implicated in insulin resistance (IVW estimate = −0.61SD (95% CI −0.95, −0.26; P=5.34×l0^−4^)) compared with a positive causal estimate when using the remaining 8 SNPs (IVW estimate = 1.26SD (95% CI 0.25, 2.27; P=0.01) (p(het) between groups <0.001) (**Figure 5**). The effect estimate obtained using the “insulin resistance” variants was largely unchanged after removing SNPs that were found to overlap with loci associated with BMI (IVW estimate = −0.53SD (95% CI −0.91, −0.15; P=0.006), although the positive causal estimate obtained using the other variants was halved (IVW estimate= 0.42SD (95% CI 0.11, 0.72; P=0.007) (**Supplementary Table 9, Supplementary Figure 6**).

**Figure 5.**
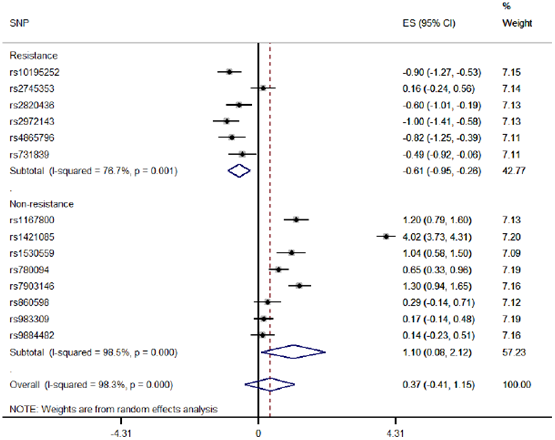
Genetic associations between BMI and fasting insulin stratified by relationship with insulin resistance

Using the more comprehensive list of “insulin resistance” variants (N=10) and “insulin secretion” variants (N=18/21 which were present in the GWAS summary results) (**Supplementary Table 10**), the effect estimates were similarly inverse when using the insulin resistance SNPs (IVW estimate = −0.72SD (95% CI −1.06, −0.38; P=4.16×l0^−5^) and positive when using “insulin secretion” variants 1.06SD (95% CI 0.34,1.79; P=0.004) (**Supplementary Table 10, Supplementary Figure 7**) although MR-Egger was less well powered to detect strong evidence for causality. Furthermore, the MR-Egger intercept was consistent with the null, providing no evidence for directional pleiotropy. However, for all stratified analyses the 12 values were very low, indicating instability in the MR-Egger estimates. For all of stratified analyses, the weighted I^2^ values were very low (0-22%), indicating instability in the MR-Egger estimates. We therefore did not implement MR-Egger regression or SIMEX adjustment for this particular analysis.

Nonetheless, for all of stratified analyses, weighted median and mode estimates were typically consistent with the IVW estimates.

## Discussion

### Summary of findings

The observational association between adiposity and insulin levels is well established^4^and the genetic correlation between the two traits has been found to be 0.65 (SE=0.06; p = 1.81×l0^−25^)^30,31^. This study used a MR approach to investigate the direction of causality in the association.

Results obtained from the analysis suggest that the positive correlation between adiposity and fasting insulin levels are at least in part explained by the causal effect of adiposity on increasing insulin. However, insufficient evidence was provided regarding a causal effect of increasing insulin on adiposity and thus the “insulin-carbohydrate” hypothesis of weight gain was not supported in this analysis.^2^

Furthermore, we identified a high degree of heterogeneity in the causal estimates obtained from the list of variants associated with insulin. This heterogeneity may be attributed to varying mechanisms of action of the insulin-associated variants. The insulin-related variants found to be inversely associated with BMI have all been determined to be “insulin resistance” SNPs in previous studies where they showed an inverse association with adiposity.^29,32^ This inverse association, which is in the opposite direction to that anticipated, has been attributed to the loci having primary effects on subcutaneous adipocyte function and distribution,^29,33,33^ although the extent of directional pleiotropy was not strong for these variants in the MR-Egger analysis.

In the stratified analysis to establish causal effects using the “insulin resistance” SNPs compared with the remaining 8 variants, there was some evidence for a positive causal estimate obtained using the remaining variants. However, this effect was halved when removing SNPs that were found to overlap with loci associated with BMI. Furthermore, as the partitioning of insulin resistance and non-resistance SNPs was determined based on their associations with HDL and triglycerides^29^, which in turn are causally influenced by BMI, this might induce spurious associations through collider bias.^34,35^ When using a more comprehensive list of “insulin secretion variants”, there was some evidence for a positive causal effect, although the Steiger test provided some evidence that the SNPs were not having a direct effect through BMI. The lack of robust causal effect is consistent with previous findings showing little evidence of an association between genetic variants associated predominantly with insulin secretion and a number of anthropometric traits,^29^ again casting doubt on the causal role of hyperinsulinemia on obesity.

### Strengths and limitations

A major strength of MR analysis in this context is that it enables an assessment of the direction of causality in an observed association between two traits. In particular, the use of genetic variants robustly associated with the exposures of interests and the two-sample design utilising publicly available data from large-scale meta-analyses have been used to maximise power to detect causal effects in this context^10,23^

This two-sample approach can also be used to address issues of weak instrument bias present in a one-sample setting which elevates the rate of Type 1 errors. In a two-sample MR analysis, any bias due to weak instruments is in the direction of the null.^36^ While this protects against inflated Type 1 error rates, this bias may lead to lower power to detect a causal effect, which could potentially explain the lack of causal effect of insulin on body mass index where the genetic instrument explained just 0.4% of the variance in insulin. However, overlap between the two datasets used in a two-sample MR approach can mitigate this bias towards the null^37^, which in the case of MAGIC and GIANT is ∼48,000 individuals.

A limitation of MR is the presence of a pleiotropic association of a genetic variant with the outcome which is independent of the exposure of interest. However, the use of MR-Egger regression, weighted median and weighted mode techniques have been used to provide estimates of causal effects that are robust to the presence of pleiotropy in different but complementary ways. While the sensitivity analyses left the causal estimate of BMI on insulin levels unchanged, the estimate from MR-Egger was larger than the IVW estimate for the causal effect of insulin on BMI. However, a limitation of the MR-Egger approach highlighted in this work was the violation of the InSIDE assumption due to the presence of the same variant in the SNP list for the exposure and the outcome. When overlapping SNPs were excluded, the causal effect of insulin on BMI was reduced in both the IVW and MR-Egger analyses.

Efforts (led by the MAGIC consortium) are already underway to identify additional variants associated with fasting insulin levels with the results of a trans-ethnic analysis involving more than 280,000 non-diabetic individuals from 144 studies currently being prepared.^38^ Preliminary results suggest 62 loci (43 novel) associated with fasting insulin. While the addition of these loci to the set available for future MR analyses may improve our ability to predict fasting insulin as an exposure (i.e. increase the variance explained), the difficulties highlighted in this analysis regarding heterogeneity of effects and potential pleiotropy are only likely to be enhanced. Additional work is needed to determine the functionality of the insulin-related variants in order to obtain the most reliable variants to improve the validity of the MR analysis assessing the causal effect of insulin on adiposity. Furthermore, while in this analysis we only assessed the causal effect of fasting insulin, the causal impact of other glycemic traits, including insulin response, on adiposity remain to be fully established.

### Implications

Overall, no clear causal effect of insulin on BMI was established compared with the strong effect of increasing BMI on insulin levels. This analysis draws to question the validity of the alternative hypothesis of weight gain based on anabolic effects of insulin induced by a high-sugar diet^2^ The lack of causal effect of insulin on body mass index is aligned with findings demonstrating no genetic link between sugar metabolism and BMI assessed using copy number variation at the human amylase locus^39^ and a genetic variant in *FGF21* recently associated with sweet intake and preference^40^.

N.B. Results based on random-effects meta-analysis so differ slightly from those based on IVW in **Supplementary Table 8**

## Acknowledgements

We would like to thank Jie Zheng for his help with the implementation of LD Hub http://ldsc.broadinstitute.org/1 to estimate genetic correlation and sample overlap.

## References

1. Hill JO. Understanding and addressing the epidemic of obesity: an energy balance perspective. Endocr Rev 2006; 27(7): 750–61.

2. Taubes G. The science of obesity: what do we really know about what makes us fat? An essay by Gary Taubes. Bmj 2013; 346: f1050.

3. Ludwig DS, Friedman MI. Increasing adiposity: consequence or cause of overeating? Jama 2014; 311(21): 2167–8.

4. Kahn SE, Hull RL, Utzschneider KM. Mechanisms linking obesity to insulin resistance and type 2 diabetes. Nature 2006; 444(7121): 840–6.

5. Cottrell RC. Essay was based on incorrect premises and an unproved assumption. Bmj 2013; 346: f3120.

6. Smith R. Are some diets “mass murder"? Bmj-Brit Med J 2014; 349.

7. Te Morenga L, Mallard S, Mann J. Dietary sugars and body weight: systematic review and meta-analyses of randomised controlled trials and cohort studies. Bmj 2013; 346: e7492.

8. Heller S. Weight gain during insulin therapy in patients with type 2 diabetes mellitus. Diabetes research and clinical practice 2004; 65 Suppl 1: S23–7.

9. Davey Smith G, Ebrahim S. 'Mendelian randomization': can genetic epidemiology contribute to understanding environmental determinants of disease? International journal of epidemiology 2003; 32(1): 1–22.

10. Davey Smith G, Hemani G. Mendelian randomization: genetic anchors for causal inference in epidemiological studies. Human molecular genetics 2014; 23(R1): R89–98.

11. Davey Smith G, Ebrahim S. What can mendelian randomisation tell us about modifiable behavioural and environmental exposures? Bmj 2005; 330(7499): 1076–9.

12. Pierce BL, Ahsan H, Vanderweele TJ. Power and instrument strength requirements for Mendelian randomization studies using multiple genetic variants. International journal of epidemiology 2011; 40(3): 740–52.

13. Burgess S, Dudbridge F, Thompson SG. Combining information on multiple instrumental variables in Mendelian randomization: comparison of allele score and summarized data methods. Stat Med 2016; 35(11): 1880–906.

14. Burgess S, Butterworth A, Thompson SG. Mendelian randomization analysis with multiple genetic variants using summarized data. Genet Epidemiol 2013; 37(7): 658–65.

15. Corbin LJ, Richmond RC, Wade KH, et al. BMI as a Modifiable Risk Factor for Type 2 Diabetes: Refining and Understanding Causal Estimates Using Mendelian Randomization. Diabetes 2016; 65(10): 3002–7.

16. Timpson NJ, Nordestgaard BG, Harbord RM, et al. C-reactive protein levels and body mass index: elucidating direction of causation through reciprocal Mendelian randomization. Int J Obes (Lond) 2011; 35(2): 300–8.

17. Bowden J, Davey Smith G, Burgess S. Mendelian randomization with invalid instruments: effect estimation and bias detection through Egger regression. International journal of epidemiology 2015; 44(2): 512–25.

18. Bowden J, Davey Smith G, Haycock PC, Burgess S. Consistent Estimation in Mendelian Randomization with Some Invalid Instruments Using a Weighted Median Estimator. Genet Epidemiol 2016; 40(4): 304–14.

19. Locke AE, Kahali B, Berndt SI, et al. Genetic studies of body mass index yield new insights for obesity biology. Nature 2015; 518(7538): 197–206.

20. Scott RA, Lagou V, Welch RP, et al. Large-scale association analyses identify new loci influencing glycemic traits and provide insight into the underlying biological pathways. Nature genetics 2012; 44(9): 991–1005.

21. Aschard H, Vilhjalmsson BJ, Joshi AD, Price AL, Kraft P. Adjusting for heritable covariates can bias effect estimates in genome-wide association studies. American journal of human genetics 2015; 96(2): 329–39.

22. Dupuis J, Langenberg C, Prokopenko I, et al. New genetic loci implicated in fasting glucose homeostasis and their impact on type 2 diabetes risk (vol 42, pg 105, 2010). Nature genetics 2010; 42(5): 464-.

23. Hemani G, Zheng J, Wade KH, et al. MR-Base: a platform for systematic causal inference across the phenome using billions of genetic associations. BiorXiv 2017.

24. Cook RD. Detection of Influential Observation in Linear-Regression. Technometrics 1977; 19(1): 15–8.

25. Bowden J, Del Greco MF, Minelli C, Davey Smith G, Sheehan NA, Thompson JR. Assessing the suitability of summary data for two-sample Mendelian randomization analyses using MR-Egger regression: the role of the I2 statistic. International journal of epidemiology 2016.

26. Hartwig FP, Davey Smith G, Bowden J. Robust inference in two-sample Mendelian randomisation via the zero modal pleiotropy assumption. BiorXiv 2017.

27. Steiger JH. Tests for Comparing Elements of a Correlation Matrix. Psychol Bull 1980; 87(2): 245–51.

28. Hemani G, Tilling K, Davey Smith G. Orienting the causal relationship between imprecisely measured traits using genetic instruments. BiorXiv 2017.

29. Scott RA, Fall T, Pasko D, et al. Common genetic variants highlight the role of insulin resistance and body fat distribution in type 2 diabetes, independent of obesity. Diabetes 2014; 63(12): 4378–87.

30. Bulik-Sullivan B, Finucane HK, Anttila V, et al. An atlas of genetic correlations across human diseases and traits. Nature genetics 2015; 47(11): 1236–41.

31. Bulik-Sullivan BK, Loh PR, Finucane HK, et al. LD Score regression distinguishes confounding from polygenicity in genome-wide association studies. Nature genetics 2015; 47(3): 291–5.

32. Kilpelainen TO, Zillikens MC, Stancakova A, et al. Genetic variation near IRS1 associates with reduced adiposity and an impaired metabolic profile. Nature genetics 2011; 43(8): 753–U58.

33. Lotta LA, Gulati P, Day FR, et al. Integrative genomic analysis implicates limited peripheral adipose storage capacity in the pathogenesis of human insulin resistance. Nature genetics 2017; 49(1): 17–26.

34. Aschard H, Vilhjalmsson BJ, Joshi AD, Price AL, Kraft P. Adjusting for Heritable Covariates Can Bias Effect Estimates in Genome-Wide Association Studies. American journal of human genetics 2015; 96(2): 329–39.

35. Day FR, Loh PR, Scott RA, Ong KK, Perry JRB. A Robust Example of Collider Bias in a Genetic Association Study. American journal of human genetics 2016; 98(2): 392–3.

36. Pierce B, Burgess S. Efficient Design for Mendelian Randomization Studies: Subsample and Two-Sample Instrumental Variable Estimators. Am j Epidemiol 2013; 177: S117–S.

37. Burgess S, Davies NM, Thompson SG. Bias due to participant overlap in two-sample Mendelian randomization. Genetic Epidemiology 2016; 40(7): 597–608.

38. Prokopenko I, Investigators ftM-AoGal-rtCM. Four glycaemic trait trans-ethnic genome-wide association meta-analyses using densely imputed genetic data in up to 281,416 non-diabetic individuals. http://wwwabstractsonlinecom/Plan/ViewAbstractaspx?sKev=0fabb0a0-c691-4714-9ea8-45cf543335ee&cKev=d373e545-ee73-40h8-hh35-0efaa0flc079&mKey=l5a3630e-7769-4d64-a80a-47fl90ac2f4f 2017.

39. Usher CL, Handsaker RE, Esko T, et al. Structural forms of the human amylase locus and their relationships to SNPs, haplotypes and obesity. Nature genetics 2015; 47(8): 921–+.

40. Soberg S, Sandholt CH, Jespersen NZ, et al. FGF21 Is a Sugar-Induced Hormone Associated with Sweet Intake and Preference in Humans. Cell Metab 2017; 25(5): 1045–+.

